# Dopamine modulates dynamic decision-making during foraging

**DOI:** 10.1101/709857

**Authors:** Campbell Le Heron, Nils Kolling, Olivia Plant, Annika Kienast, Rebecca Janska, Yuen-Siang Ang, Sean Fallon, Masud Husain, Matthew A J Apps

## Abstract

The mesolimbic dopaminergic system exerts a crucial influence on incentive processing. However, the contribution of dopamine in dynamic, ecological situations where reward rates vary, and decisions evolve over time, remains unclear. In such circumstances, current (foreground) reward accrual needs to be compared continuously with potential rewards that could be obtained by travelling elsewhere (background reward rate), in order to determine the opportunity cost of staying versus leaving. We hypothesised that dopamine specifically modulates the influence of background – but not foreground – reward information when making a dynamic comparison of these variables for optimal behaviour. On a novel foraging task based on an ecological account of animal behaviour (marginal value theorem), human participants were required to decide when to leave locations in situations where foreground rewards depleted at different rates, either in rich or poor environments with high or low background rates. In line with theoretical accounts, people’s decisions to move from current locations were independently modulated by both foreground and background reward rates. Pharmacological manipulation of dopamine D2 receptor activity using the agonist cabergoline significantly affected decisions to move on, specifically modulating the effect of background but not foreground rewards rates. In particular, when on cabergoline, people left patches in poor environments much earlier. These results demonstrate a role of dopamine in signalling the opportunity cost of rewards, not value per se. Using this ecologically derived framework we uncover a specific mechanism by which D2 dopamine receptor activity modulates decision-making when foreground and background reward rates are dynamically compared.

**Significance statement:** Many decisions, across economic, political and social spheres, involve choices to “leave”. Such decisions depend on a continuous comparison of a current location’s value, with that of other locations you could move on to. However, how the brain makes such decisions is poorly understood. Here, we developed a computerized task, based around theories of how animals make decisions to move on when foraging for food. Healthy human participants had to decide when to leave collecting financial rewards in a location, and travel to collect rewards elsewhere. Using a pharmacological manipulation, we show that the activity of dopamine in the brain modulates decisions to move on, with people valuing other locations differently depending on their dopaminergic state.

## INTRODUCTION

The mesolimbic dopaminergic system plays a crucial role in motivating behaviour towards goals and has been closely linked to neural circuits which convey information about incentives (Schultz and Dickinson, 2000; Haber and Knutson, 2010; Salamone and Correa, 2012; Manohar et al., 2015; Hamid et al., 2016; Le Bouc et al., 2016). Several experiments across species have demonstrated a crucial role for dopamine in overcoming costs to obtain rewards (Salamone and Correa, 2012; Le Bouc et al., 2016; Syed et al., 2016; Le Heron et al., 2018b) and for learning about rewarding outcomes to update future behaviour (Pessiglione et al., 2006; Schultz, 2016). Tasks probing dopamine function typically require an agent to make *binary decisions* between presented options, based on learning the contingent relationship between stimuli and rewards, or an integration of cost and reward information (Salamone et al., 2007; Schultz, 2016; Le Heron et al., 2018b). Yet, in real world settings many of our decisions are not binary choices between stimuli or actions. Moreover, animal models increasingly highlight that dopamine signals change gradually during ongoing behaviours as the *rate* of obtaining rewards changes, suggesting a need to examine the role of dopamine in decisions that are dynamic in nature (Howe et al., 2013; Hamid et al., 2016; Mohebi et al., 2019).

One real-world, dynamic decision, is whether to stay in a current location or switch to an alternative to maximize reward collection (Pearson et al., 2014; Mobbs et al., 2018). Such decision-making requires a continuous comparison between the current (foreground) reward rate and the alternative (background) reward rate available in the environment (Rutledge et al., 2009; Kurniawan et al., 2011; Constantino et al., 2017). However, despite the clear ecological significance of such reward rate comparisons for decisions to move on, dopamine’s role in modulating these processes remains unclear.

It has been proposed that tonic (slower-changing) dopamine signals encode information about environmental richness, and background reward rates (Niv et al., 2007). This is supported by voltammetry experiments linking slow changes in dopamine levels to a rodent’s reward environment (Hamid et al., 2016), and evidence of changes in motor vigour as dopamine state varies in humans (Beierholm et al., 2013; Guitart-Masip et al., 2014; Le Bouc et al., 2016; Le Heron et al., 2018b). However this association has been questioned (Zenon et al., 2016), and it remains unknown whether the link between dopamine and background reward rates applies to more abstract – but ecologically crucial – decisions to move on.

Models of foraging behaviour, derived in behavioural ecology, provide an ideal theoretical framework to investigate the relationship between dopamine and dynamic human decision-making. Marginal Value Theorem (MVT), an influential foraging model, provides a formal framework for how animals decide to leave a location (“patch”) as rewards deplete, and travel to find rewards in another (Charnov, 1976; Stephens and Krebs, 1986). At its core is the notion that animals should continuously compare the instantaneous foreground reward rate with the average background reward rate, and an optimal forager should leave when the former falls below the latter (Charnov, 1976; Stephens and Krebs, 1986; Pearson et al., 2014). Within MVT, these two rates independently impact when it is optimal to leave patches, making this dynamic decision framework ideal for testing if dopamine processes the background reward rate. However, despite the behaviour of a wide range of species following MVT predictions (Nonacs, 2001; Stephens et al., 2007), little is known about whether such principles extend to human behaviour (Hutchinson et al., 2008; Pearson et al., 2014; Mobbs et al., 2018; Gabay and Apps, 2019).

We developed a novel ecologically derived decision-making task in which participants chose when to move on as foreground and background reward rates varied. We hypothesised that both young and old human participants would make leaving decisions in accordance with MVT, and that manipulating dopamine receptor activity using the D2-agonist cabergoline would selectively modulate the influence of *background reward rate* on decisions to move on.

## MATERIALS AND METHODS

To test the hypothesis that manipulating dopamine availability would modulate the influence of background reward rates on patch-leaving decisions in humans we designed a novel foraging based task. In the first study, we highlight the validity of this task in healthy young participants. In the second, we manipulated dopamine availability pharmacologcally, testing the influence of cabergoline administration on older adults in double-blind, placebo-matched, crossover design.

### Participants

This study was approved by the local research ethics committee and written informed consent was obtained from all participants.

#### Study ONE

40 healthy volunteers (mean age 24, range 20-30) were recruited via a local database. One was subsequently excluded because of poor engagement with the task (identified at de-briefing).

#### Study TWO

30 healthy older (mean age 69, range 60-78) participants were recruited via a local database. Potential participants were screened for the presence of neurological, psychiatric or cardiovascular diseases, or for the use of medications that could interact with cabergoline, and excluded if any of these were present. One subject was subsequently excluded because a core metric of task performance (variance in leaving times per condition) fell outside three standard deviations of the mean variance, leaving 29 participants for analysis.

### Experimental design

All participants were administered a computer based patch-leaving task in which they had to decide when to move on from a current patch. The task design independently manipulated background and foreground reward rates, based on the principles of MVT, a theory of optimal foraging behaviour (Charnov, 1976; Stephens and Krebs, 1986). The task was framed as a farming game in which people had to collect as much milk (reward) as possible – this would be sold at a market at the end of the game and their financial remuneration was according to the milk accrued. Participants spent a fixed time in each of two farms, collecting milk from fields of cows and making decisions of whether to move on (leave the current field for the next one) (**Figure 1A**). Moving on to the next field incurred a time cost (travel time) during which no milk could be collected.

**Figure 1.**
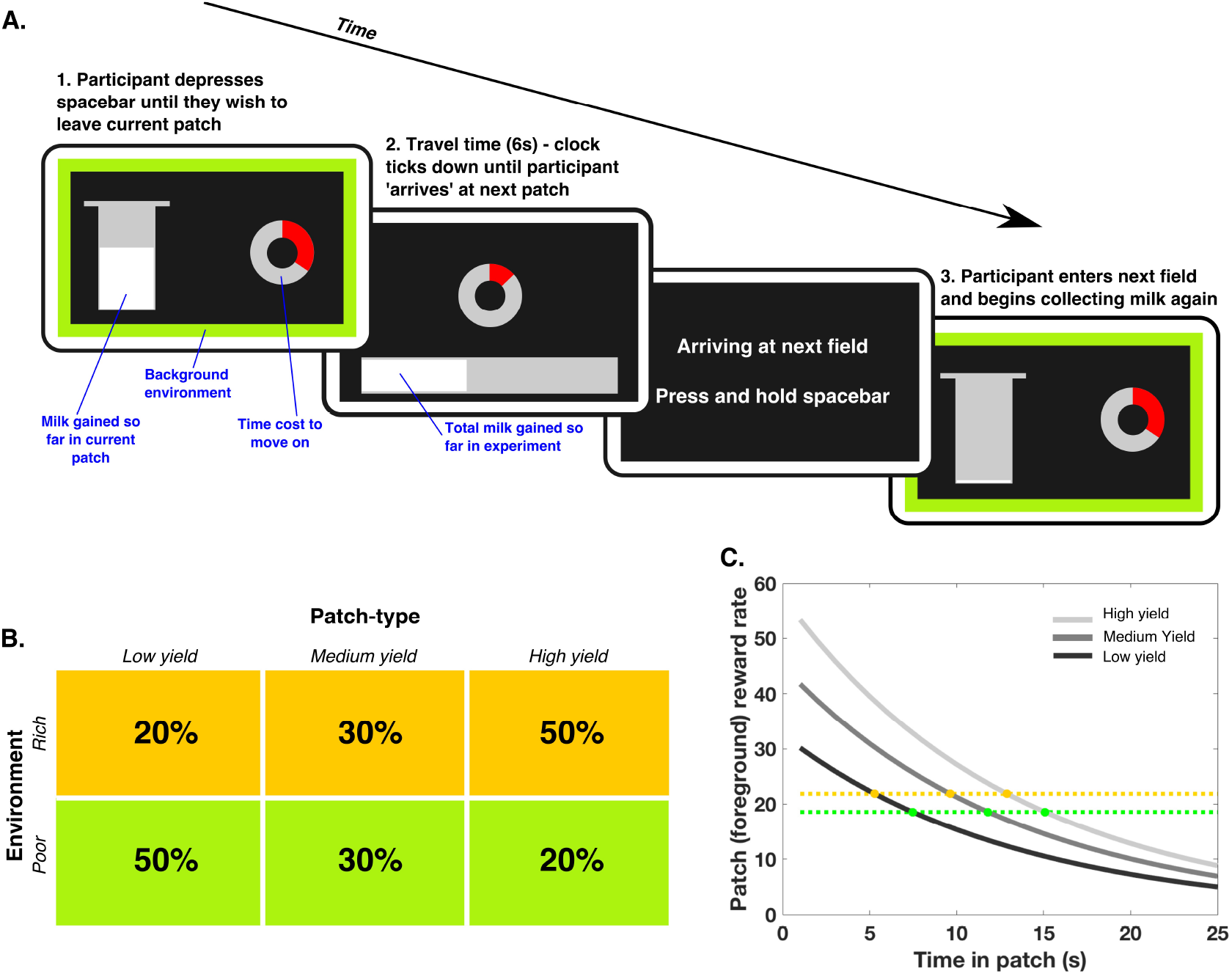
Patch leaving paradigm. (**A**) Participants had to decide how long to remain in their current patch (field), in which reward (milk) was returned at an exponentially decreasing rate (displayed on the screen by continuous filling (white bar) of the silver bucket), before moving on to the next patch, which incurred a fixed cost of 6 seconds during which they could collect no reward. Their goal was to maximise milk return across the whole experiment. The instantaneous rate of bucket filling indicated the **foreground reward rate**, whilst the coloured frame indicated the distribution of different patch types, and thus the **background reward rate**. Participants were aware they had approximately 10 minutes in each environment, but were not shown any cues to indicate how much total time had elapsed. Following a leave decision, a clock ticking down the 6 second travel time was presented. (**B**) Three foreground patch-types were used, differing in the scale of filling of the milk bucket (low, medium and high yield), which determined the foreground reward rate. Two different background environments (farms) were used, with the background reward rate determined by the relative proportions of these patch-types. The rich environment contained a higher proportion of high yield fields, and a lower proportion of low yield ones, meaning it had a higher background reward rate than the green farm, which had a higher proportion of low yield fields. (**C**) According to MVT participants should leave each patch when the instantaneous reward rate in that patch (grey lines) drops to the background environmental average (gold and green dotted lines). Therefore, people should leave sooner from all patches in rich (gold dotted line) compared to poor (green dotted line) environments, but later in high yield compared to low yield patches. Crucially, these two effects are independent from each other.

Participants aimed to maximise their overall reward returns by deciding how long to spend in these sequentially encountered patches, in which the current (foreground) reward rate decreased in an exponential manner. The reward obtained so far in the patch was displayed as a bucket which continuously filled during patch residency.

Three patch-types were used, differing in the scaling factor of the reward function (***S*** in equation one below), and corresponding to low (32.5), medium (45) and high (57.5) yield patches. The foreground reward rate, after ***T*** seconds in a patch, was determined by the equation:

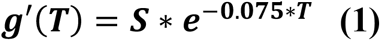

The height of milk displayed in the bucket was proportional to the integral of equation (1) between time = 0 and T, and was updated with a frequency of 20Hz. Participants were not explicitly instructed which patch-type they were currently in – rather they inferred this by observing the rate of milk accumulation.

The background reward rate was manipulated by varying the proportions of low, medium and high yield patches in “farms”, in a pseudorandomised fashion (**Figure 1B**). In the rich farm (environment), 50% of the patches were high yield, 30% medium and 20% low yield, whilst in the poor farm 50% of the patches were low yield, 30% medium and 20% high. Therefore, the background reward rate was higher in the rich environment. The background reward rate was continuously cued by the coloured border on the screen, indicating either the rich (gold border) or poor farm (green border). MVT demonstrates that, to maximise reward gain, participants should leave each field when the instantaneous reward rate in the field (from **equation 1**) drops below the background average reward rate for the farm. Simply, for a given patch-type, participants should leave earlier in the rich environment compared to the poor environment (**Figure 1C**).

### Procedure

Before commencing the experiment participants were trained on the task elements using a structured explanation and practice session lasting ~20 minutes. Comprehension of the different elements was checked verbally before commencing the main experiment, with volunteers asked to explain what each display item meant. They were not given any instructions as to what optimal behaviour would be. However, they were told they would spend an equal amount of time on the two farm types (gold and green) and that they would never run out of fields. Participants were seated in front of a desktop computer running Pyschtoolbox (pyschtoolbox.org) implemented within MATLAB (MathWorks, USA).

When participants chose to leave their current patch (by releasing the spacebar they had been holding down), they incurred a fixed time cost of 6 seconds, described as the time to walk to the next patch. During this time a counter was displayed which ticked down the seconds until the next patch was reached. On arriving at the next patch participants were cued to “press and hold the spacebar”, and after doing this the screen display changed to show the new patch

#### Study ONE

Participants were tested in a single session following training as above.

#### Study TWO

This was conducted as a randomised, double-blind, placebo-controlled study. Participants were tested in two separate sessions, once following administration of a single dose of 1mg cabergoline (which stimulates post-synaptic D2 receptors (Brooks et al., 1998)) and once following administration of an indistinguishable placebo tablet. An older population was chosen because they may have a relative dopaminergic deficit compared to younger people (Karrer et al., 2017) and thus be more sensitive to the intervention (Fallon et al., 2019). The order of testing was counterbalanced across drug manipulation, gender and order of background foraging environment (rich-poor or poor-rich).

### Statistical analyses

We used a hierarchical linear mixed effects model (*fitlme* in MATLAB, Mathworks, USA; maximum likelihood estimation method) as our primary analysis method for both experiments, to account for between and within subject effects. All fixed effects of interest (patch, environment and where applicable dopamine) and their interactions were included, and the random effects structure was determined by systematically adding components until the Akaike Information Criterion was minimised (Barr et al., 2013). Notably the signifncance of any effects in all these models were the same simpler models fitting only a random effect of subject. Significant model effects were also followed up with parametric tests (t-tests and/or Analysis of Variance).

#### Study ONE

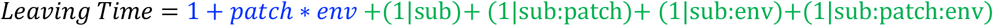

#### Study TWO

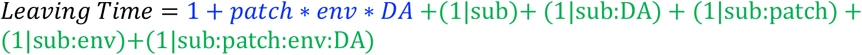

*patch = foreground reward rate, env = background reward rate, DA = dopamine state, sub = subject. Fixed effects are shown in blue, random effects in green.*

To avoid the potentially biasing effects of outlying data points on the primary analysis we excluded, subject by subject, any trials in which the leaving time was more than 3 standard deviations from that individual’s mean leaving time. Of note, this approach did not change the significance (or otherwise) of any reported results compared to analysis of the full data set.

## RESULTS

### Healthy human foragers are guided by MVT principles

Within MVT, foreground and background reward rates should have independent effects on how long an individual remains in a patch. People should leave low yield patches sooner than high yield patches, and patches in rich environments sooner than patches in poor environments. In line with these hypotheses, in Study 1, we found a main effect of foreground reward, as well as a main effect of background reward, but no interaction on participants’ (N = 39) decisions about when to leave their current patch (Foreground: F(1,74.6) = 528, p < 0.0001; Background: F(1,37.5) = 40, p < 0.0001; Foreground × Background: F(1,1929) = 1.6, p = 0.2; **Table 1A**). Furthermore, behaviour conformed to predicted directionality of these effects, with higher patch yield, and poor compared to rich background environment, both leading to later patch leaving times (**Figure 2A & 2B**).

**Table 1.**
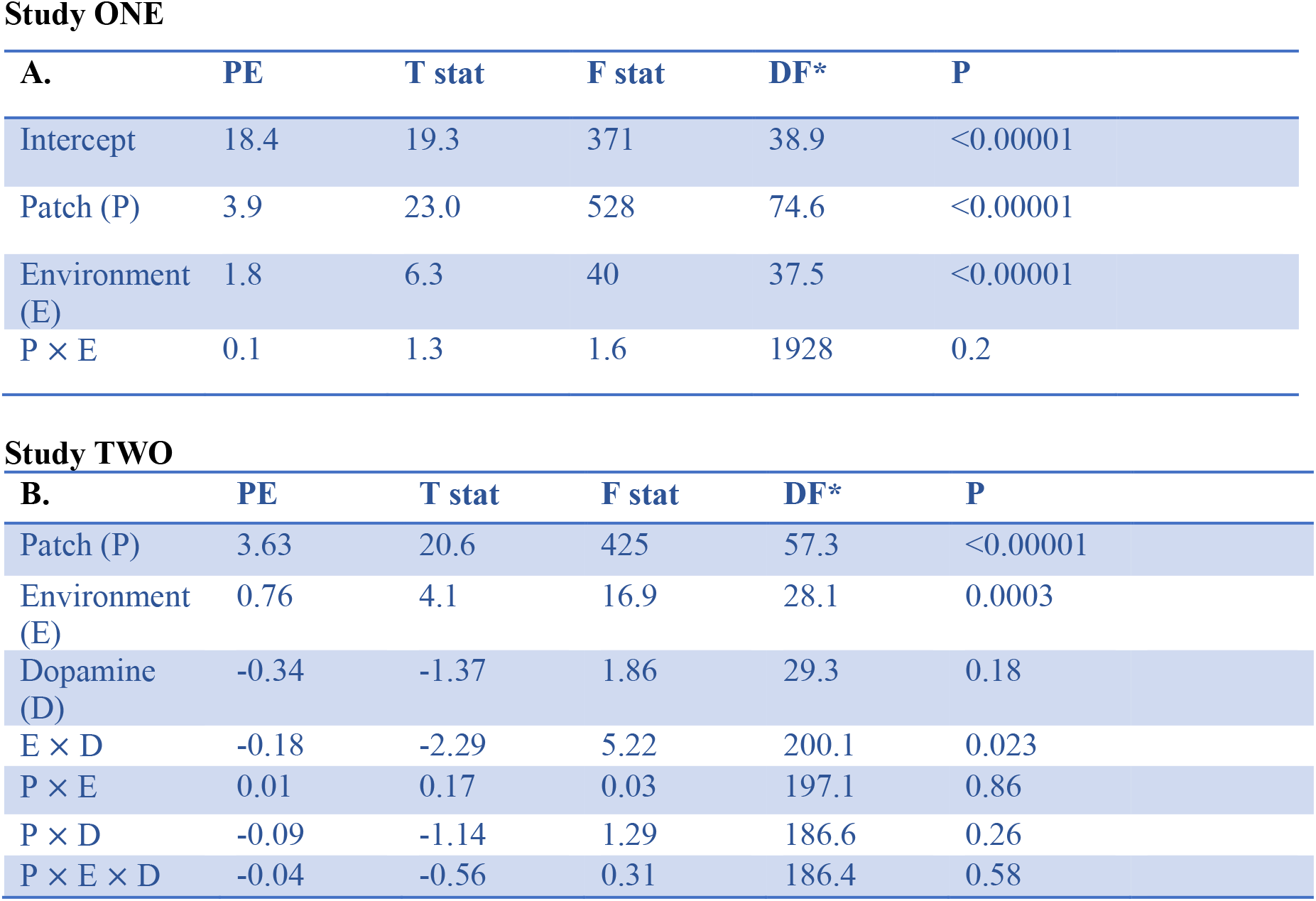
Linear mixed effects models from each experiment. * DF were calculated using the Satterthwaite correction method.

Study ONE
| A. | PE | T stat | F stat | DF* | P |
| --- | --- | --- | --- | --- | --- |
| Intercept | 18.4 | 19.3 | 371 | 38.9 | <0.00001 |
| Patch (P) | 3.9 | 23.0 | 528 | 74.6 | <0.00001 |
| Environment (E) | 1.8 | 6.3 | 40 | 37.5 | <0.00001 |
| P × E | 0.1 | 1.3 | 1.6 | 1928 | 0.2 |

**Table 1. Linear mixed effects models from each experiment**
| B. | PE | T stat | F stat | DF* | P |
| --- | --- | --- | --- | --- | --- |
| Patch (P) | 3.63 | 20.6 | 425 | 57.3 | <0.00001 |
| Environment (E) | 0.76 | 4.1 | 16.9 | 28.1 | 0.0003 |
| Dopamine (D) | -0.34 | -1.37 | 1.86 | 29.3 | 0.18 |
| E × D | -0.18 | -2.29 | 5.22 | 200.1 | 0.023 |
| P × E | 0.01 | 0.17 | 0.03 | 197.1 | 0.86 |
| P × D | -0.09 | -1.14 | 1.29 | 186.6 | 0.26 |
| P × E × D | -0.04 | -0.56 | 0.31 | 186.4 | 0.58 |

**Figure 2.**
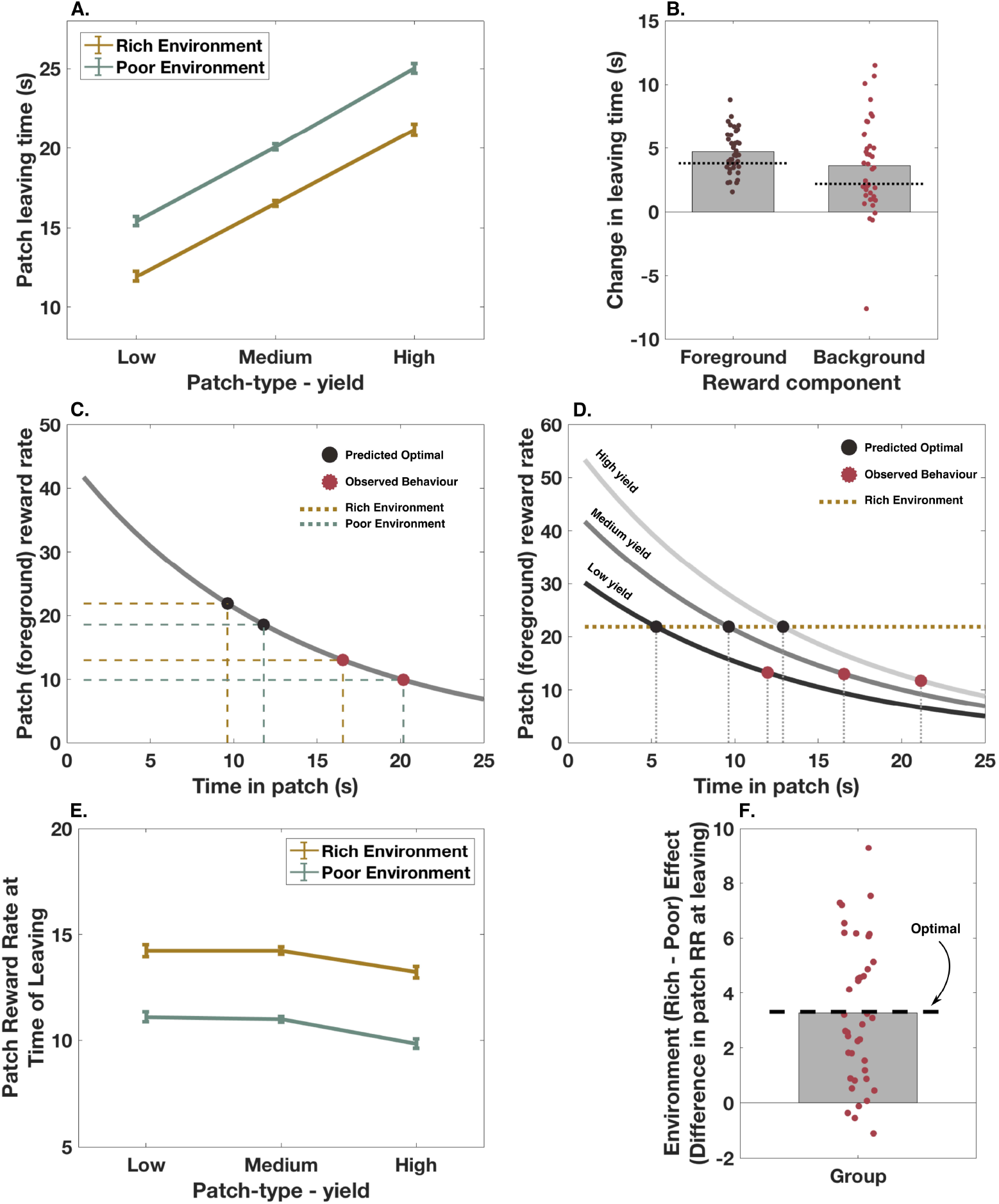
Healthy human foragers are guided by MVT principles. (**A**) Raw patch leaving times. Participants (N = 39) left patches later when the background environment was poor, compared to rich (p < 0.00001), and when patches had higher, compared to lower yields (p < 0.00001), with no interaction between patch-type and background environment (p = 0.2). (**B**) These effects of changing reward parameters were in the predicted direction, with participants leaving on average 4.7s later as patch-type varied, and 3.6s later in poor compared to rich environments. There was more variation between individuals in the effects of changing background, compared to foreground, reward rates. Dashed lines show predicted (MVT) effects of changing reward rate on leaving time. **(C) & (D)** Participants showed a bias to remain in patches longer than predicted by MVT. Mean leaving time for each environment, collapsed across patch-type is shown in (C), whilst (D) demonstrates mean leaving times for each patch-type in the rich environment. (**E**) The foreground (patch) reward rate at which participants chose to leave each patch varied as a function of background environmental richness (rich vs poor). (**F**) The magnitude of this background environment effect was close to optimal (as predicted by MVT). Error bars are ± SEM.

### Are healthy people optimal foragers?

Although participants showed effects in the directions predicted by MVT, we wanted to know whether the *magnitude* of these effects conform to foraging theories, which stipulate the optimal time to leave each patch. Every individual showed a significant bias to *remain longer* across all patch types (across both environments) than optimal, on average leaving 8.0s later than MVT predictions (t_38_ = 8.4, p < 0.001**, Figure 2C & 2D)**. However, it has been noted that non-human primates also show such a bias to stay, but are close to optimal once controlling for this bias, e.g., by analysing the *relative changes* across conditions (Hayden et al., 2011). Therefore for each participant we subtracted their own mean leaving time from each of their patch leaving decisions, and calculated the magnitude of the *background* (poor-rich) and *foreground* (mean change between each patch-type) reward rate effects (**Figure 2B & 2F**).

MVT makes two core predictions about behaviour as foreground and background reward rates change, which can be used to assess optimality of foraging behaviour (independent to any systematic bias to remain in patches longer). Firstly, in the background environments (poor vs rich), the foreground reward rate at leaving a given patch-type should differ by the same amount. Secondly, foragers should adjust their leaving time as patch quality varies, such that the instantaneous reward at leaving is the same in each patch (for a given background). That is within an environment, each patch should be left, regardless of its yield, when the rate at which milk is being accrued is the same.

Strikingly, participants varied their leaving times as background environment changed, such that the difference in reward rate between the two conditions was not significantly different from the predicted optimal difference (mean difference in reward rate at leaving = 3.33, actual difference between environments if behaving optimally = 3.30, t_38_ = 0.07, p = 0.95, **Figure 2E & 2F**). In contrast, the foreground reward rate at patch leaving did vary across patch type (F(1.5,42) = 6.73, p = 0.005, RM-ANOVA). Although the instantaneous reward rate on leaving low and medium yield patches did not differ (mean difference = 0.06, t_37_ = 0.2, p = 1), participants remained in high yield patches until the instantaneous reward rate was lower compared to both medium yield (mean difference = 1.1, t_37_ = 3.8, p = 0.002), and low yield patches (mean difference = 1.1, t_37_ = 2.6, p = 0.04; **Figure 2E**).

In summary, across participants’ sensitivity to changes in the foreground were not quite optimal, but on average participants adjusted leaving times in response to changes in the background environment in a manner that closely matched the actual changes in background reward rate. They also adjusted their leaving behaviour such that the reward rate at leaving did not differ between low and medium yield patches, although they tended to leave high yield patches later (i.e., after patch reward rate had dropped further). Thus, human behaviour on this task broadly conformed to MVT principles, and was close to being optimally influenced by background and foreground reward rates, although people were not precisely optimal. This is despite no instructions of what pattern of behaviour would maximise rewards in the task.

### Cabergoline alters the use of background reward information to guide patch leaving

Having demonstrated that healthy human patch leaving behaviour is aligned with the predictions of MVT, particularly in response to changes in background reward rate, we next examined whether dopamine modulates the effect of background reward rate (environment) on patch leaving behaviour. Using a within-subjects design, in Study 2, leaving times for 29 healthy older people on placebo or following administration of the D2 receptor agonist cabergoline were analysed using a LME model.

Firstly, the main effects reported in Study 1 were replicated. Both foreground (patch) and background (environment) reward rates significantly influenced patch leaving time, and there was no interaction between the two (Foreground: F(1,57) = 425, p < 0.0001; Background: F(1,28) = 16.9, p = 0.0003; Foreground x Background: F(1,197) = 0.03, p = 0.86; **Figure 3A and 3B**). Furthermore, the magnitude of effect of both background and foreground reward rate on leaving time did not significantly differ between the young and older groups [mean difference (young – old) Foreground = 0.2s, t_66_ = 0.49, p = 0.62; mean difference (young – old) Background = 1.2s, t_66_ = 1.77, p = 0.08].

**Fig 3.**
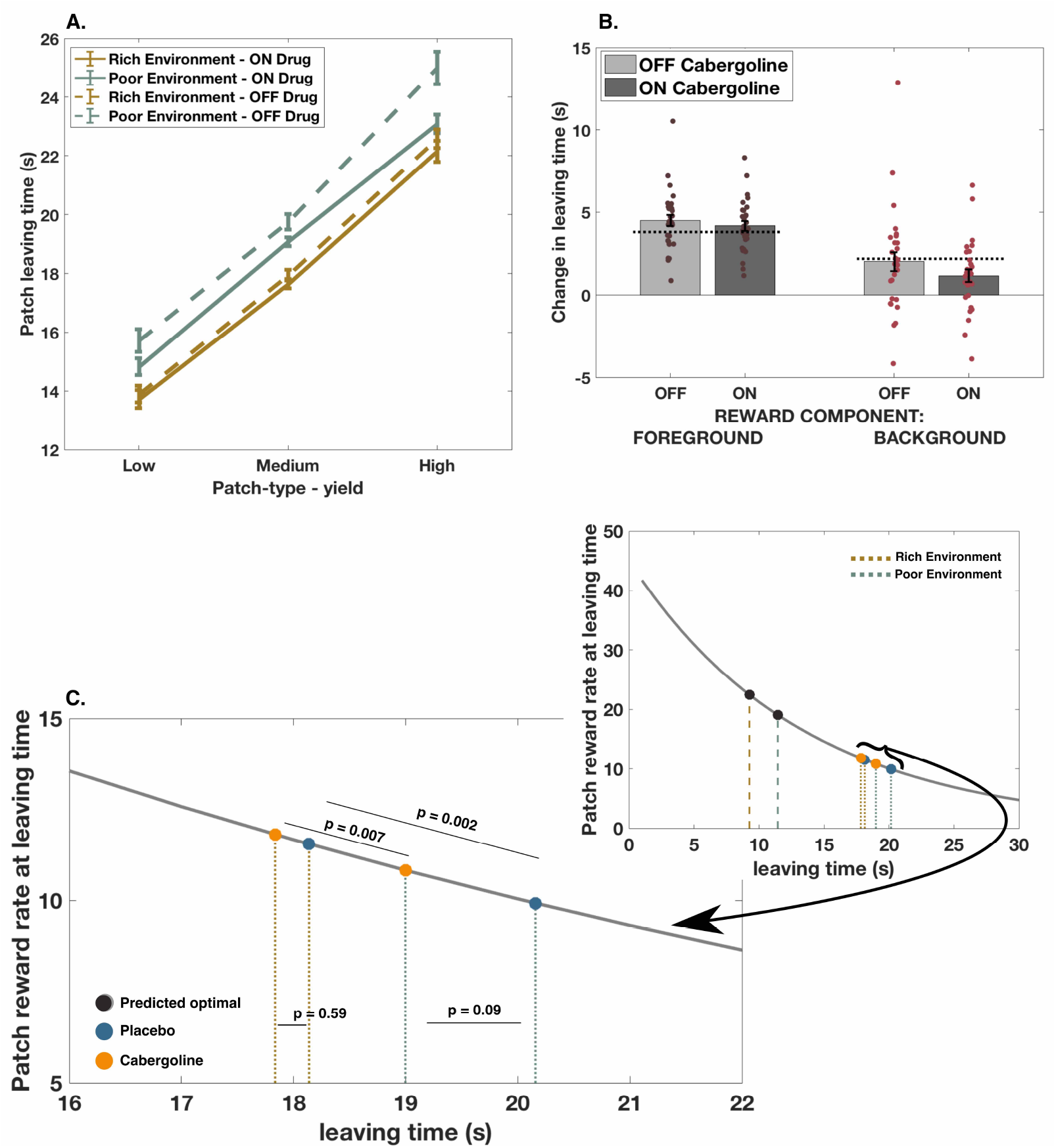
Cabergoline alters use of background reward information to guide patch leaving. (**A**) Mean patch leaving times for each patch-type, split by environment and drug state. **(B)** There was a significant interaction between drug and background (environment) reward rate on leaving time, with a reduced effect of background environment ON cabergoline compared to OFF (p = 0.023*). Green dotted lines in (A) and represent the predicted magnitude of effect of the manipulation, based on the marginal value theorem*. (B) In contrast, there was no significant interaction between drug and the effect of foreground (patch) reward rate on patch leaving (p = 0.26). (**C**) Raw leaving times for the two groups, in rich (gold) and poor (green) environments (collapsed across patch-type). The reduced effect of drug seemed mainly driven by participants ON cabergoline leaving patches earlier – and therefore when the patch reward rate was higher – in poor environments. (**D**) There was no main effect of cabergoline on either the instantaneous reward rate at patch leaving, or on raw leaving times (not shown) when collapsing across environments. N = 29, comparisons are within-subject, error bars are ± SEM.

There was a significant interaction between drug state and the effect of background reward rate on leaving time (F(1,200) = 5.22, p = 0.023, **Table 1B**). When ON cabergoline, people were less sensitive to the difference between poor and rich environments than when OFF drug even though they still showed a significant effect of background environment both ON and OFF the drug (**Figure 3A & 3B)**. Post-hoc analysis suggests this interaction was driven by people leaving patches in the poor environment much earlier ON cabergoline than OFF, but only leaving patches in the rich environment slightly earlier ON compared to OFF (mean difference (OFF – ON) poor environment = 1.2s, t_28_ = 1.75, p = 0.009; mean difference (OFF – ON) rich environment = 0.3s, t_28_ = 0.59, p = 0.56; **Figure 3C**).

We hypothesised that modulating dopamine levels would *not* alter the effect of foreground reward rate on patch leaving, if manipulating tonic levels predominantly affects the processing of average reward rates. In line with this hypothesis, there was no significant drug × patch interaction (F(1,187) = 1.29, p = 0.26): cabergoline did *not* lead to a significant change in the way participants used foreground reward rate information to guide leaving decisions (**Figure 3B**). There was also no statistically significant difference in leaving times overall on drug compared to placebo (mean difference = 0.73s, F(1,29) = 1.86, p = 0.18), nor did the reward rate at leaving vary as a function of drug state (mean difference = 0.39, t_28_ = 0.8, p = 0.41).

As would also be predicted within MVT, there was no interaction in leaving times between foreground and background reward rate. Moreover, the observed drug × background reward rate interaction was present across all patch types, with no 3-way interaction (F(1,186) = 0.31, p = 0.58). All of these results remain significant after controlling for weight, height and BMI. Although the experiment was designed to minimise the effects of any learning, because the dopaminergic manipulations could in theory lead to differential learning effects between states we analysed the data from experiment two for session or order effects. The inclusion of session (1^st^ or 2^nd^) worsened model fit (change in BIC 7.6), and the parameter estimate for session effect was not significant (PE= −0.16, F(1,29) = 0.36, p = 0.56). Similarly, including order (the session * drug interaction) also worsened model fit (change in BIC 14.2) and again this term was not significant (PE= 0.87, F(1,29) = 1.4, p = 0.25). Therefore session and order effects were not included in the final model. The inclusion of these effects did not change the significance (or otherwise) of the other model terms.

Could participants be paying less attention when off medication? We analysed leaving time variability to examine whether participants’ decisions were more noisy as a function of drug state. There was no significant difference in the variance of each participant’s decisions between placebo and cabergoline conditions (Mean Difference _**PLAC-CAB**_ = 0.31, t_28_ = 1.34, p = 0.19). Therefore cabergoline had a specific rather than general effect on patch leaving behaviour, altering only the influence of background reward rate on leaving time.

## DISCUSSION

When to move on and leave a specific rewarding activity or location is an essential decision problem for animals and humans alike. Here, we show that humans – both young and old – make dynamic foraging decisions that, although not optimal, broadly conform to ecological principles captured by Marginal Value Theorem (MVT) (Charnov, 1976; Stephens and Krebs, 1986). Furthermore, dopaminergic D2 receptor activity may play a crucial role in modulating such decisions. Specifically, the findings support the view that dopamine plays an important role in signalling the average value of alternative locations, influencing dynamic decisions of when to move on. Administration of cabergoline altered the effect of *background* – but not *foreground* – reward rate on patch leaving times. In particular, this interaction between cabergoline and background reward rate was driven mainly by people leaving all patches in poor environments earlier.

The results provide new evidence for the role of dopamine in decision-making. Manipulation of dopamine levels modulated the influence of background reward rate on dynamic decisions about when to switch behaviour. Specifically, ON cabergoline people tended to leave *all patch-types* in the poor environment *earlier* than when OFF drug. In contrast, in the rich environment, there was a much smaller change in leaving times between the ON and OFF drug states. The drug manipulations used here putatively alter tonic dopamine levels (Brooks et al., 1998), a component of the dopaminergic neuromodulatory system which has been ascribed, in the context of motor responses, a role in signalling background reward rates (Niv et al., 2007; Hamid et al., 2016). Of course, in this study we were not able to measure firing rates of dopamine neurons. Nevertheless, some existing evidence suggests tonic dopamine levels encode information about background reward rate, and therefore the opportunity cost (alternatives that are foregone) of chosen actions (Niv et al., 2007; Guitart-Masip et al., 2014). However, others have argued that they encode a more specific signal for the value of a current action, independent of environmental context (Zenon et al., 2016). In addition, it has been argued that tonic dopamine signals the value of exploring alternative options.

It is important to appreciate though that much of the previous research in this area has used binary choices, which may not always reflect real-world problems. Furthermore, in many experiments the value of exploring or exploiting are directly opposed and have an instrumentally predictive value of obtaining an immediate (foreground) reward (Daw et al., 2006; Kayser et al., 2015; Westbrook and Frank, 2018). However, in ecological settings, choices to “leave” a patch and explore are not choices between two stimuli with a predictive value, but instead involve travelling to obtain rewards elsewhere, and the rewards available in a patch can be orthogonal to the environment one is in. Using a novel experimental paradigm, based on theories of animal foraging, we show the effect of D2 activity specifically on background reward rate, in a potentially more ecologically important decision process. These findings suggest dopamine is at the core of how foraging decisions are made. Furthermore, the specificity of cabergoline for D2 receptors suggests D2-mediated pathways may be of particular importance for signalling such contextual reward information (Beaulieu and Gainetdinov, 2011).

What mechanisms might drive such changes? Recently it has been suggested that foraging problems share parallels with evidence accumulation, where the foreground reward rate serves as the evidence accumulating towards a decision bound, or threshold, set by the background environment. When the foreground rate reaches that threshold it triggers a decision to leave (Davidson and El Hady, 2019). Our results suggest that dopamine signals carry information about *when* to leave, by setting a threshold for the foreground reward rate, that is dependent on the richness of the environment. Thus, patches should be left in richer environments sooner, and at a higher foreground reward-rate, than in poor environments because less evidence – or a higher reward rate – is needed in order to leave a patch. Such effects are consistent with the absence of an interaction between foreground and background reward rates that we find in healthy young participants, that is also a key principle of MVT. When D2 receptors were stimulated (ON state), people left patches earlier (at a higher foreground reward rate) in the poor environment, consistent with an increase in perceived richness of the environment. Thus, whilst the average reward rate may increase the vigour of movements or exploratory binary choices, in more abstract, ecological decision settings it serves to increase the perceived environmental richness, setting a higher threshold reward rate of when to leave.

Importantly, these results appear to be driven by changes in sensitivity to the background reward rate, rather than alternative explanations., Firstly, we used a continuously changing patch gain function where rewards were constantly accrued – rather than stepped changes as has been used in previous studies (Hutchinson et al., 2008; Constantino and Daw, 2015). This approach has the advantage of minimising the use of simple heuristics to guide decisions while having the statistical advantage of leaving the dependent variable approximately normally distributed. Secondly, variance in patch leaving times did not change as a function of drug state. This makes it unlikely the results can be explained by a confounding factor such as reduced attention. Thirdly, as participants were explicitly informed of the current environment in which they were in, and had experienced the different background reward rates in a training phase, it is unlikely that such effects could be explained by differences in learning as a function of drug.

Our results highlight that human behaviour in an ecologically-derived decision-making task is closely described by a model based on the principles of MVT (Charnov, 1976; Pearson et al., 2014). This accords with earlier field work in behavioural ecology (Stephens and Krebs, 1986; Pearson et al., 2014) and anthropology (Smith et al., 1983; Metcalfe and Barlow, 1992) literatures, and more recent work beginning to explore the neural basis of such decisions (Hayden et al., 2011). In the current study, the use of a foraging framework informed by MVT enabled us to dissociate the effects of reward rates on different time scales, in a way that is not possible in reinforcement-learning based manipulations of average reward rates (Niv et al., 2007; Mobbs et al., 2018). In most reinforcement-based tasks examining foreground/background reward rates, the two are not independent. Firstly, receipt of an instrumental reward instantaneously increases average reward rate. Secondly, average reward rate influences the probability of receiving an instantaneous reward. In contrast, In a patch-leaving framework, the current foreground rate – the current value of a patch - is always declining, independent of the environment, and reward accrued in the current patch does not inform a participant about the background rate. Thus, the study design allows independent comparison of the effects of dopamine modulation on two different components of reward, compared to those used in reinforcement-learning based tasks.

Grounding the experimental design within the framework of MVT also allowed for a direct comparison of behaviour against optimal predictions, that have previously shown to hold in animals both freely foraging in the wild, and within controlled experimental setups (Krebs et al., 1977; Hayden et al., 2011). Here, participants utilised the dissociable aspects of their reward environment to adjust patch leaving behaviour in close to optimal fashion. This provides evidence for a common decision principle guiding foraging-style behaviour in both humans and other animals, and allows further investigation of the specific neural mechanisms underlying it. It highlights the importance of considering reward – not as a single construct – but rather as a multi-dimensional reinforcer, with distinct effects occurring across different time scales, underpinned by dissociable neural and neuromodulatory mechanisms (Pearson et al., 2014).

From a clinical perspective these findings may be significant when considering mechanisms underlying common disorders of motivated behaviour, such as apathy (Le Heron et al., 2019). Apathy is often associated with disruption of mesolimbic dopaminergic systems (Santangelo et al., 2015), and, at least in some cases can be improved with D2/D3 receptor agonists (Adam et al., 2013; Thobois et al., 2013). Accumulating evidence demonstrates altered reward processing in patients with apathy (Strauss et al., 2014; Le Heron et al., 2018a), and it is plausible – although as yet untested – that chronic underestimation of background environment reward leads to a state where it is never “worth switching” from a current activity, even if this activity is very minimal. Future work could profitably explore this hypothesis.

Recent theoretical accounts of decision-making have called for a shift to more ecologically derived experiments to investigate the mechanisms of this fundamental neural process (Pearson et al., 2014; Mobbs et al., 2018). The current results highlight the utility of such an approach, demonstrating a role for D2 activity in signalling the average background reward rate during foraging. This demonstrates the applicability of a model validated in wild and experimental animal populations to human behaviour. It links basic ecological models of animal behaviour to a mechanistic understanding of human decision making, highlighting the specific influence of dopaminergic systems as people decide when to move on as they pursue rewards in their environment.

## ACKNOWLEDGEMENTS

This research was supported by a University of Oxford Christopher Welch Scholarship in Biological Sciences, a University of Oxford Clarendon Scholarship and a Green Templeton College Partnership award (C.L.H.); a Wellcome Trust Principal research fellowship to MH; the NIHR Oxford BRC (Biomedical Research Centre; MH); the Velux Foundation (MH); A BBSRC David Phillips Fellowship (BB/R010668/1) to MAJA.

## AUTHOR CONTRIBUTIONS

CLH, NK, MH and MAJA designed the study; CLH, NK and MAJA coded the experiment, CLH, OP, AK, RJ and YA collected data; CLH, NK, SF and MAJA analysed data; CLH, NK, SF, MH and MAJA wrote the paper.

